# Co-expression of recombinant human collagen α1(III) chain with viral prolyl 4-Hydroxylase in *P. pastoris* GS115

**DOI:** 10.1101/2022.04.13.488258

**Authors:** Jiayuan Fang, Ze Ma, Dongyue Liu, Zhaoguo Wang, Shuo Zheng, Hongyan Wu, Peijun Xia, Xi Chen, Rui Yang, Linlin Hao

**Affiliations:** College of Animal Science, Jilin University, 5333 Xi’an Road, Changchun, Jilin 130062, China; Guoda New Materials Co., LTD, 3999 Air Street, Changchun, Jilin 130062, China

**Keywords:** hydroxylation, purification, recombinant human collagen α1(Ш) chain, virus prolyl 4-hydroxylase

## Abstract

Prolyl 4-hydroxylase (P4H) is essential to maintain the stable triple-helix structure and function of human collagen α1(Ш) chain (COL3A1). To obtain hydroxylated human COL3A1, the human *COL3A1* and the viral *P4H* A085R were co-expressed in *P. pastoris* GS115. The sequence of human *COL3A1* without N-terminal and C-terminal was selected for expression. Colony PCR analysis and sequencing after transfection showed that the target gene had inserted successfully. Real-time quantitative PCR (RT-qPCR) indicated that human *COL3A1* and *P4H* were expressed at the mRNA levels. SDS-PAGE and Western blotting analysis of supernatant from the recombinant methylotrophic yest culture showed that recombinant human COL3A1 (rhCOL3A1) was secreted into the culture medium with an apparent molecular mass of approximately 130 kDa. It was noted that the rhCOL3A1 expession quantity was higest at 120 h of induction. Furthermore, mass spectrometry analysis demonstrated that the rhCOL3A1 was expressed successfully. His-tagged rhCOL3A1 protein was purified by Ni-affinity column.

## 1. Introduction

Collagens are the main structural protein of vertebrates, widely distributed in connective tissues of animals, and are also one of the most abundant protein families in the human body. Collagens play the indispensable role in diverse biological functions, such as maintaining normal physiological functions, repairing injuries, and effecting cell attachment and proliferation (Gelse et al. 2003). There are many types of collagens, all of which composed of triple-helical structure, in which COL3A1 is the key to supporting the epidermis (Vesentini et al. 2005). The application of COL3A1 is wide, mainly in sponge dressings (Rao 1995), membrane dressings, gel dressings and other medical fields. Nowadays, COL3A1 used in these fields is mainly from connective tissue of animal, such as porcine, bovine, asinine and piscine skin. However, it needs to be considered that animal collagen extracted from bone and buffalo skin by traditional methods has certain safety risks. For this reason, expressing recombinant collagen by genetic engineering with low cost and high efficiency has received increasing attention (Olsen et al. 2003).

During the synthesis of collagen, hydroxyproline plays an important role in maintaining the triple-helix structure and biological properties of collagen (Shoulders and Raines 2009). And many organisms, such as mammalian cells with P4H, can directly form hydroxylated collagen by post-translational modification (Geddis and Prockop 1993; Fichard et al. 1997). However, due to the lack of prolyl hydroxylase, some prokaryotic expression systems, like bacteria, cannot hydroxylate proline residues in collagen (Lamberg et al. 1996). In general, human collagen is coexpressed with P4H to achieve hydroxylation modification (Pinkas et al. 2011; Xu et al. 2011; Nokelainen et al. 2001). Human P4H is an α2β2 heterotetramer, and its limited stability and complex structure may limit efficient production of recombinant hydroxylated collagen in bacterial and yeast populations lacking endogenous hydroxylase (Neubauer et al. 2007; Winter and Page 2000).

Recently, some non-human P4Hs were of concern. Several researchers have demonstrated that the co-expression of P4H α and PDI/β genes in the *Escherichia coli* (*E*.*coli*) could produce active 4-hydroxylase (Neubauer et al. 2005). And some plant-derived P4H showed substrate preference (Kaska et al. 1987). Recombinant hydroxylated collagen was obtained by co-expression with proline hydroxylase L593 from giant mimicry virus in *E*.*coli* (Rutschmann et al. 2014). Moreover, it has been confirmed that recombinant hydroxylated collagen can be co-expressed with collagen in mammalian cells, plant cell cultures and yeast (Xu et al. 2011). Besides, a strain of P4H virus isolated from *Paramecium bursaria Chlorella* virus-1 (PBCV-1) with 242-residue polypeptide was similar to the α subunit sequence of human P4H. The substrates of this P4H were not restricted (Eriksson et al. 1999). In addition, the methylotrophic yeast *P. pastoris* has proved to be an effective production system for the production of heterologous proteins (Baez et al. 2005).

At present, rhCOL3A1 is mostly expressed in small fragments, because its full-length expression requires the hydroxylation of proline hydroxylase. However, the commonly used bacterial expression system and yeast expression system do not contain proline hydroxylase, and the stability of the product obtained by the recombinant expression is very low (Lamberg et al. 1996). In order to obtain effective and stable rhCOL3A1, the gene encoding for human *COL3A1* without N-terminus and C-terminus was constructed into pPIC9K expression vector, and the viral *P4H* A085R derived from PBCV-1 was constructed into pPICZA expression vector, then the two vectors were co-expressed in *P*.*pastoris* GS115 to obtain hydroxylated collagen. The expression vectors of *COL3A1* and *P4H* were successfully constructed and secreted rhCOL3A1 was obtained.

## 2. Materials and methods

### 2.1. Strains, Plasmids

*E*.*coli* strain DH5 (ANGYU, Shanghai, China) was used to amplify plasmids. *P. pastoris* GS115 strain (ANGYU) was used as the host to express the heterologous protein. The expression vectors pPIC9K and pPICZA were procured from Invitrogen.

### 2.2. Construction of rhCOL3A1 expression plasmid

The expression vector of the human COL3A1 was constructed as follows: First, the cDNA sequence of the core functional region of human *COL3A1* (GenBank: BC028178.1) was amplified by PCR with primers designed. Next, the product was digested with *SnaBI*-*NotI* and cloned into expression vector pPIC9K at the same sites to obtain the pPIC9K-COL3A1 plasmid. Besides, the sequence of human *COL3A1* was without N-terminus and C-terminus but with the α-factor signal sequence. The expression vector of human *P4H* was constructed as follows: the cDNA sequence of sub A085R protein gene sequence(GenBank:NP-048433.1) was selected as proline hydroxylase gene sequence (Lu et al. 1995). Next, the gene encoding P4H was amplified by PCR, and the product was digested with *EcoRI*-*KpnI* and cloned into expression vector pPICZA at the same sites resulting in the pPICZA-P4H plasmid.

### 2.3. Transformation

Plasmid pPIC9K-COL3A1 and plasmid pPICZA-P4H were linearized at the site with sacI before electrotransformation. After mixing 20 μl linearized recombinant plasmids with 80 μl competent *P. pastoris* GS115 cells, the cells were immediately transferred to a prechilled electroporation cuvette and incubated on ice for 15 min. The charging voltage was 1.5 kV, capacitance was 25 mF, and resistance was 200Ω. The transformed plasmids were coated on yeast extract peptone dextrose medium (YPD) containing bleomycin resistance, and the culture plates were incubated at 30°C for 2∼3 days to grow.

### 2.4. PCR analysis of integrants

To further confirm the integration of pPIC9K-COL3A1 and pPICZA-P4H plasmids into the genome of *P*.*pastoris* GS115 growing on YPD containing bleomycin resistance plates, genomic PCR was performed. Briefly, the clones growing on YPD plates were expanded in the liquid YPD medium, and their genomic DNA was isolated using TIANamp Yeast DNA Kit (TIANGEN, Beijing, China). Two pairs of primers for PCR were 5’/3’*AOX1* and α-factor/3’ *AOX1*, and the sequence of 5’*AOX1* was 5’-GACTGGTTCCAATTGACAAGC-3’, 3’*AOX1* was 5’-GCAAATGGCATTCTGACATCC-3’ and *α-factor* was 5’-TACTATTGCCAGCATTGCTGC-3’.

The integration of pPIC9K-COL3A1 gene was verified by PCR using 300 ng genomic DNA as template. For amplification controls, 100 ng of recombinant plasmids pPIC9K-COL3A1 and pPICZA-P4H were used as positive controls; 100 ng of empty plasmids pPIC9K were used as negative control; 100 ng of plasmids pPIC9K-COL3A1 were used as single plasmid control.

### 2.5. Shake-Flask cultivation

The successfully transfected colonies were selected for protein induction expression analysis. Single colony was cultured in buffered glycerol medium [BMGY(0.2% biotin, 1% yeast nitrogen base, 1% glycerol, 1% yeast extract, 2% peptone, 100 mM potassium phosphate)] at 30 °C with vigorous shaking (200 rpm) for 24 h. The colonies were harvested by centrifugation at 5,000 *g* for 20 min at 4 °C, and then the cell pellets were resuspended in buffered methanol medium [BMMY(0.2% biotin, 1% methanol, 1% yeast nitrogen base, 1% yesat extract, 2% peptone, 100 mM potassium phosphate)] for rhCOL3A protein induction. Methanol was supplemented every 24 h to maintain 1% methanol concentration, and the culture lasted for 144 h. After induction, the culture supernatant was collected by centrifugation at 5,000 *g* for 20 min at 4 °C.

### 2.6. RNA extraction and RT-qPCR

Total RNA was isolated using TRIzol Reagent (Takara, USA) and total RNA was harvested at different times after induction. The RNA was immediately used by the Thermo Scientific Revert Aid First Strand cDNA Synthesis Kit(Thermo, Waltham, MA, USA) for reverse transcription reactions. The qPCR reactions were carried out in a 96-well plate in CFX96 Real-time PCR Detection System (Bio-Rad). The primers were listed in Table 1. All samples were performed in triplicate. Each mRNA quantitative data represented the average of at least three measurements. The Ct value of GAPDH gene was normalized to the Ct value of target gene. Data analysis was performaed using the following formula: 2^-△△Ct^(△Ct=Ct_COL3A1/P4H_-Ct_GAPDH_)

**Table 1.**
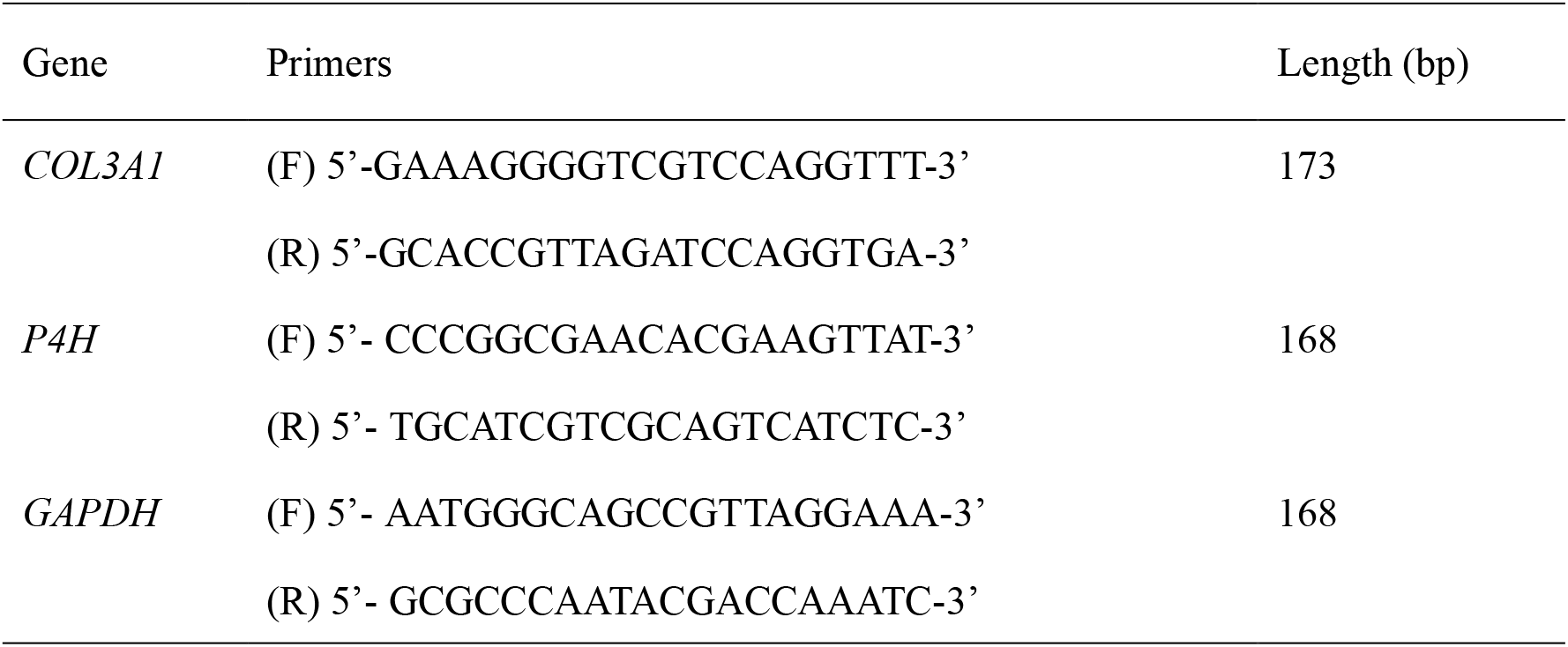
Primer sequence of *COL3A1, P4H* and *GAPDH*.

### 2.7. SDS-PAGE and Western Blotting

The expression of rhCOL3A1 was evaluated by SDS-PAGE analysis of microbial supernatant with 12% Tris-glycine gels by the method of Laemmli(Laemmli 1970). The protein sizes on the gel was measured by approximate comparison with the color-mixed Protein Marker (Solarbio, Beijing, China). Then, the 8% Tris-glycine gels containing proteins were transferred to polyvinylidene fluoride (PVDF) membranes. After blocking with western blocking buffer at 25 °C, the first memberane incubated with anti-COL3A1 mouse monoclonal antibody (Santa Cruz, USA) at 1:100 dilution overnight at 4 °C. The second memberane incubated with His-Tag Monoclonal antibody (Proteintech, USA) at 1:3000 dilution overnight at 4 °C. Then the two membranes were washed with TBST three times (5 min each), and the first memberane was incubated with goat anti-mouse IgG labeled with horseradish peroxidase (TIANGEN) at 1:1000 dilution at 25 °C for 2 h. BeyoECL Plus (Beyotime, Shanghai, China) assessed immune reactivity.

### 2.8. Mass spectrometry analysis

After the recombinant protein was isolated by SDS-PAGE, the 130 kDa protein band of rhCOL3A1 was cut from gels and sent it to the Sangon Biotech for Liquid chromatography coupled with tandem mass spectrometry (LC-MS/MS) analysis.

### 2.9. Purification

The yeast fermentation medium was harvested after 120 h culture. Medium supernatant was first clarified by centrifugation at 5,000 *g* for 20 min at 4 °C. Filtering the centrifuged supernatant with a 0.4 μm filter, then were directly loaded onto a Ni-NTA 6 FF Pre-Packed Gravity column (Sangon Biotech, Shanghai, China) preequilibrated with 10 × Binging Buffer. The column was washed with Elution Buffer and collected 4 mL protein. The purified rhCOL3A1 was analyzed by SDS-PAGE with 8% Tris–glycine gels, and protein ladder was used as reference. The concentration of rhCOL3A1 was determined by the BCA protein.

### 2.10. Statistical Analysis

All experiments were performed in triplicate. All statistical analyses were performed using GraphPad Prism 9.0 (GraphPad Software, San Diego, CA). Differences were considered significant at p < 0.05.

## 3. Results and Discussion

### 3.1. Design of COL3A1 sequence and Construction of vector

Many studies have demonstrated that recombinant collagen can be obtained by genetic engineering (Olsen et al. 2005; Luther et al. 2011), and rhCOL3A1 had the same properties as human COL3A1. In this study, we cloned and co-expressed the *P4H* and human *COL3A1* in *P*.*pastoris* GS115. The cDNA sequence of COL3A1 core functional region (GenBank: BC028178.1) was not contain N-terminus and C-terminus, which were replaced by α-MF signal peptide sequence and six histidine coding sequences, respectively (Li et al. 2015). The *P4H* gene sequence was obtained from viral A085R protein gene sequence(LU. et al. 1995). According to the codon preference of *P. pastoris* cells, *COL3A1* and *P4H* gene sequences were optimized for expression in *P. pastoris* GS115.

High copy number *P. pastoris* expression vectors pPIC9K and pPICZA were selected. Vector pPIC9K was digestion with *SnaBI*-*NotI* and vector pPICZA was digestion with *EcoRI*-*KpnI*. Fig. 1A and B showed the construction of pPIC9K-COL3A1 and pPICZA-P4H plasmids. DNA sequencing results suggested that the target genes were successfully constructed into the expression vector (data not shown). Then, the constructed plasmid would be transfected into *P. pastoris* GS115 expression host.

**Fig. 1.**
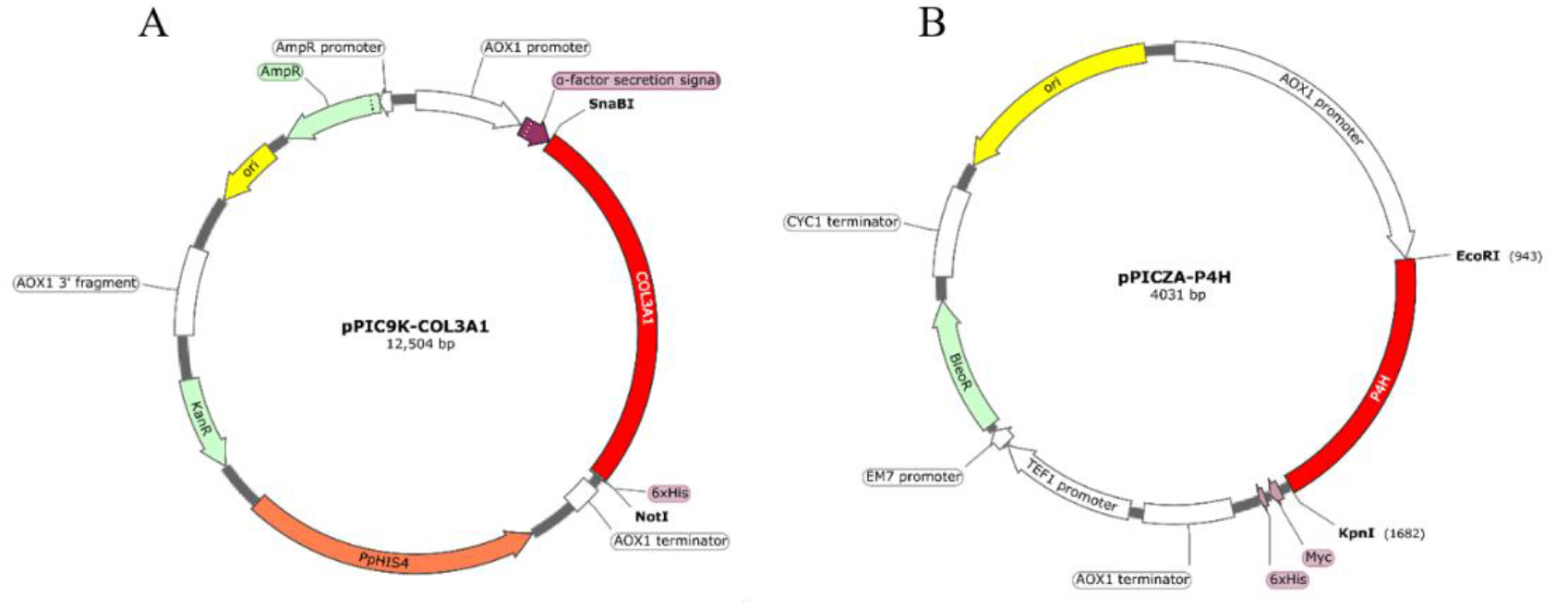
Physcial maps of the vectors. The red areas were target genes. (A) Schematic map of the recombinant expression vector pPIC9K-COL3A1. (B) Schematic map of the recombinant expression vector pPICZA-P4H.

### 3.2. Colony identification and RT-qPCR

To screen the GS115 colonies integrated rh*COL3A1* gene, Colonies PCR were applied to select recombinant strains(Xu et al. 2015), and two primer pairs were designed, respectively. The results showed that the recombinant plasmids were successfully transferred into *P*.*pastoris* GS115 (Fig. 2A). The result of CP1 and CP2 colonies PCR using 5’/3’*AOX1* primers had two bands at 2250 bp and 741 bp, respectively. The results of CP1 and CP2 colonis PCR using α –F /3’ primers had a band at 3994 bp respectively. The results were in accord with expectations.Then the single colony Cp1 and Cp2 on the [1.0 mg/ml] zeocin plate were picked for induction culture. The induced strains were collected every 24 h and RNA was extracted for RT-qPCR (Xu et al. 2015). Fig. 2B and C showed that RT-qPCR results of time-course of *P4H*. The results suggested that the expression of *P4H* gene in CP1 and CP2 strains was the highest at 96h. Fig. 2D and E showed that RT-qPCR results of time-course of *COL3A1*. The mRNA relative expression levels of *COL3A1* gene was highest at 120 h. The results might confirm that the expression of P4H can stabilize the structure of COL3A1, and this result needed to be confirmed by further experiments. Besides, the study of Vuorela et al. had shown that the co-expression of *P4H* and *COL3A1* not only maintains the stability of COL3A1 triple helix structure, but also markedly increased the amount of active prolyl 4-hydroxylase tetramer (Vuorela et al. 1997).

**Fig. 2.**
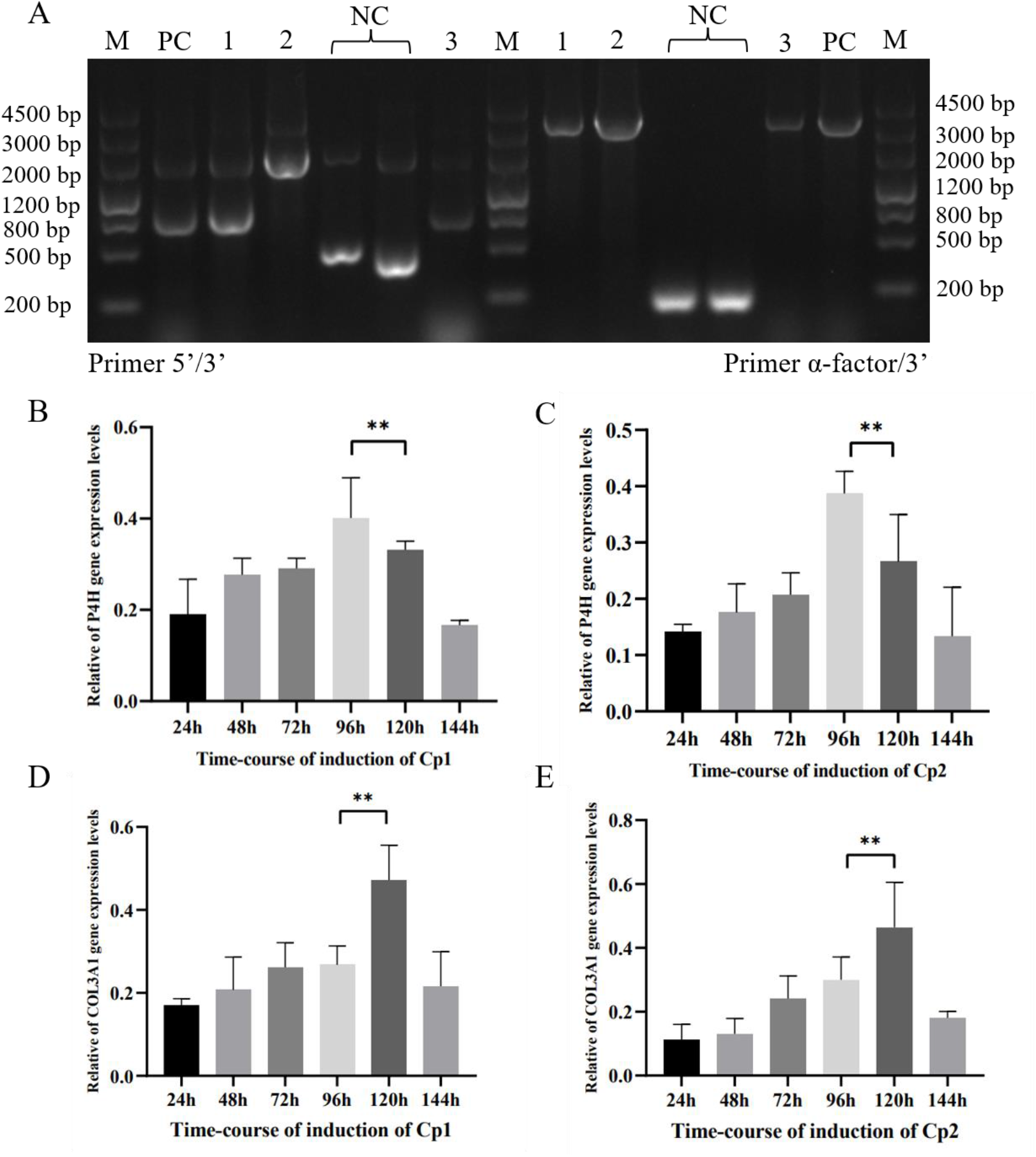
(A) Colony PCR results. Lane M, DNA ladder; Lane PC, positive control; Lane 1-3, Cp1, single plasmid (pPIC9K-COL3A1) control and Cp2; Lane NC, negative control. (B) and (C) The relative of P4H expression levels on mRNA. (D) and (E) The relative of COL3A1 expression levels on mRNA.

### 3.3. Protein identification and Purification

To confirm the rhCOL3A1 protein, SDS-PAGE and western blotting were applied. The Fig.3A was the SDS-PAGE analysis of extracelluar proteins. SDS-PAGE showed a thin protein at 130 kDa (marked in Fig.3A), and western blotting showed that the apparent molecular weight of protein was approximately 130 kDa (marked in Fig.3B), and this result was similar to the studies results of Li et al. (Li et al. 2015). Therefore, we can confirm that the *COL3A1* was expressed in the *P*.*pastoris* GS115. Besides, Fig.3B showed the rhCOL3A1 protein levels of two time-courses, and the results demonstrated that the expression levels of protein was highest for 120 h. Fig.3C suggested the P4H protein expression levels for 96 and 120 h of NC and Cp2, and the 96 h was higher, which were consistent with the qRT-PCR results (Fig.2C). Studies have shown that increasing the copy number of expressing plasmid could increase the production of recombinant protein in *P. pastoris* (Clare et al. 1991; Sreekrishna et al. 1997). Then, the rhCOL3A1 proteins were purified by Ni-affinity column and gel filtration chromatography (Shi et al. 2017). The results suggested that the rhCOL3A1 was concentrated in the first 2 ml of the eluent (Fig.3D), and whether the first 2 ml contains rhCOL3A1 fragments was something we needed to further verify in the futureand. Thus, rhCOL3A1 could be successfully purified with his-tagged.

**Fig. 3.**
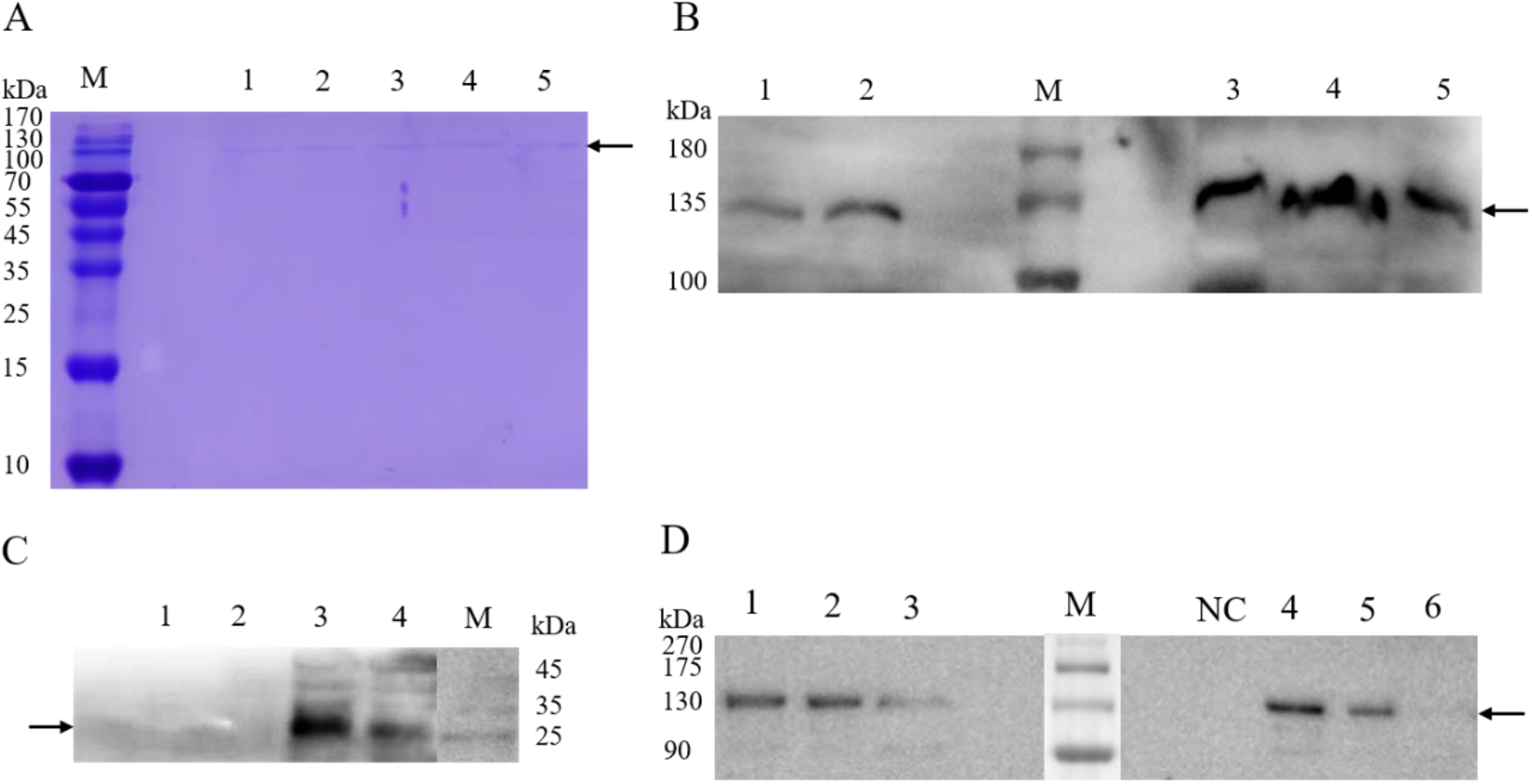
SDS-PAGE and Western blotting results for rhCOL3A1 expression. (A) SDS-PAGE results for rhCOL3A1. (B) Western blotting results for rhCOL3A1. Lane M was the standard molecular weights. Lanes 1 and lane 2 were the secretory proteins of the Cp1 strain at different induction times: 96 and 120 h, respectively, after induction. Lane 3, lane 4 and lane 5 were the secretory proteins of the Cp2 strain at different induction times: 96, 120 and 144 h, respectively, after induction. (C) Western blotting results for P4H. Lane M was the standard molecular weights. Lane 1 and lane 2 were the endocrine proteins of the negative control strain at different induction times: 96 and 120 h. Lane 3 and lane 4 were the endocrine proteins of the Cp2 strain at different induction times: 96 and 120 h. (D) Western blotting results for purified rhCOL3A1. NC stands for negative control. Lane M was the standard molecular weights. Lanes 1, lane 2 and lane 3 were the collected proteins of the Cp1 strain at different stage: 1st milliliter, 2nd milliliter and 3rd milliliter, respectively. Lanes 4, lane 5 and lane 6 were the collected proteins of the Cp2 strain at different stage: 1st milliliter, 2nd milliliter and 3rd milliliter.

### 3.4. LC-MS/MS results

LC-MS/MS was becoming an increasingly important analytical technology in the identification of peptides and proteins (Brun et al. 2009; Kirkpatrick et al. 2005). Besides, Lund et al. identified lysine methylation sites in globle proteins by mass spectrometry (Lund et al. 2019). At the same time, Li et al. directly detected mercaptan modification of various proteins in pancreatic β cells by mass spectrometry (Li et al. 2021). To confirm the extent of the proline hydroxylation in the rhCOL3A1 protein, the gel pieces containing the 130 kDa protein were cut and identified by LC-MS/MS. Fig.4A-F were the secondary spectrum of peptides identified by mass spectrometry. The abscissa was m/z and the ordinate was relative intensity. The peaks in the figure were signal peaks of fragment ions, and the peaks marked in color were matched fragment ions. It was generally believed that the more fragments ions, the more sufficient the identification evidence. The results showed that the sequence coverage was 19%, because the concentration of protein samples was low. Then, compared with UniProt proteins (rhCOL3A1 and P02461), the protein was identified as human collagen α1(III) chain (Fig.4G). Among the amino acids detected in the peptide sequence was hydroxyprolines, as shown in Fig.4H that mark in gray represented the hydroxylated amino acids. The results suggested that the proline was hydroxylated and its structure was maintained by the formation of intramolecular hydrogen bonds (Berisio et al. 2004). In summary, P4H was the key enzyme in collagen synthesis (Myllyharju 2003). The results confirmed that virus-derived P4H could play the same function as human P4H in Pichia pastoris cells.

**Fig. 4.**
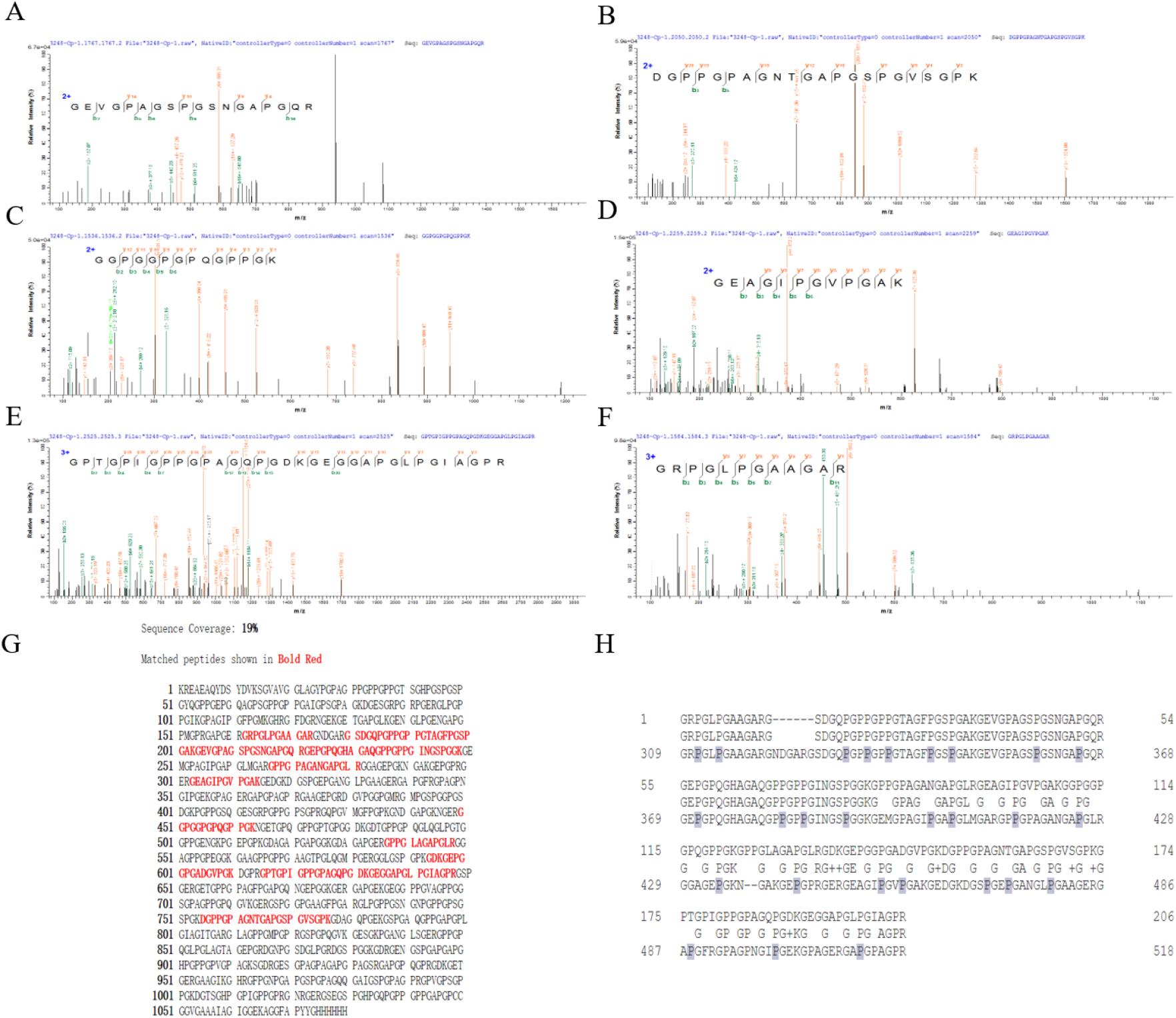
The identification of rhCOL3A1 by LC-MS/MS. (A-F) Secondary mass spectrometry of peptides. The abscissa of the secondary mass spectrometry was m/z, and the ordinate was the relative intensity. The peaks were the signal peak of the fragment ions, and the peaks marked in color were the matched fragment ions. (G) Mascot search results showed that the sequence coverage was 19%, and matched peptides was shown in bold red. (H) The hydroxylated amino acids marked in gray.

## 4. Conclusions

In this study, we successfully constructed the eukaryotic expression system of rhCOL3A1, and also have shown that the heterologous protein rhCOL3A1 with molecular weight up to 130 kDa could be secret by *P. pastoris* GS115. The results indicate that *P. pastoris* GS115 was a good host for rhCOL3A1 protein expression. The results of RT-qPCR suggested that high expression of *P4H* can maintain the stability of COL3A1. Besides, SDS-PAGE and western blotting results showed that the protein expression level of strain was the highest after 120 h of induction, consistent with the RT-qPCR results. At the same time, we confirmed that rhCOL3A1 was modified by P4H, and the presence of hydroxyproline residues was verified by LC-MS/MS analysis. In summary, we confirmed the co-expression of rhCOL3A1 with viral P4H in *P. pastoris* GS115. The strategy provided an alternative for production and purification of the rhCOL3A1 protein.

## Data Availability Statement

The original contributions presented in the study are includedin the article/supplementary material, further inquiries can bedirected to the corresponding author

## Acknowledgments

We thank members of the Hao laboratory for suggestions. LH, JF and ZM designed the experiments. JF, ZM and DL performed the experiments with assistance of ZW, SZ, HW, PX, XC and RY. JF, ZM and LH analyzed the data. JF, ZM, DL and LH wrote the manuscript.

